# Impulsive choice in mice lacking paternal expression of *Grb10* suggests intra-genomic conflict in behavior

**DOI:** 10.1101/281873

**Authors:** Claire L. Dent, Trevor Humby, Katie Lewis, Andrew Ward, Reiner Fischer-Colbrie, Lawrence S. Wilkinson, Jon F Wilkins, Anthony R. Isles

**Affiliations:** Behavioural Genetics Group, MRC Centre for Neuropsychiatric Genetics and Genomics, Neuroscience and Mental Health Research Institute, Cardiff University, Hadyn Ellis Building, Maindy Road, Cardiff, United Kingdom; Behavioural Genetics Group, School of Psychology, Cardiff University, Tower Building, Cardiff, United Kingdom; Department of Biology and Biochemistry, University of Bath, Bath, United Kingdom; Department of Pharmacology, Innsbruck Medical University, Innsbruck, Austria; Ronin Institute, Montclair, NJ 07043, USA

**Keywords:** genomic imprinting, Nesp55, delayed reinforcement, intra-genomic conflict, Grb10

## Abstract

Imprinted genes are expressed from one parental allele only as a consequence of epigenetic events that take place in the mammalian germ line and are thought to have evolved through intra-genomic conflict between parental alleles. We demonstrate, for the first time, oppositional effects of imprinted genes on brain and behavior. Specifically, here we show that mice lacking paternal *Grb10* make fewer impulsive choices, with no dissociable effects on a separate measure of impulsive action. Taken together with previous work showing that mice lacking maternal Nesp55 make more impulsive choices this suggests that impulsive choice behavior is a substrate for the action of genomic imprinting. Moreover, the contrasting effect of these two genes suggests impulsive choices are subject to intra-genomic conflict and that maternal and paternal interests pull this behavior in opposite directions. Finally, these data may also indicate that an imbalance in expression of imprinted genes contributes to pathological conditions such as gambling and drug addiction, where impulsive behavior becomes maladaptive.

## INTRODUCTION

Imprinted genes are expressed from one parental allele only as a consequence of epigenetic events that take place in the mammalian germ line (Ferguson-Smith 2011). Functionally, imprinted genes converge on specific biological processes that have prominent importance in mammals, such as *in utero* growth, metabolism and behavior (Peters 2014; Wilkins *et al.* 2016). Many of the effects that maternally-and paternally-expressed imprinted genes have on *in utero* growth and during the early post-natal period are oppositional in direction (Haig and Graham 1991; Madon-Simon *et al.* 2014), lending support to the theory that imprinting has evolved as a consequence of intra-genomic conflict (Moore and Haig 1991; Wilkins and Haig 2003).

*Grb10* is currently a unique example of an imprinted gene that is expressed from the maternal allele only in some tissues, and the paternal allele only in others (Arnaud *et al.* 2003; Hikichi *et al.* 2003; Monk *et al.* 2009). Consequently, different parental alleles of *Grb10* mediate distinct physiological functions (Garfield *et al.* 2011). Maternal *Grb10* is expressed in many peripheral body tissues, whilst being excluded from the central nervous system (Garfield *et al.* 2011), and plays a prominent role in controlling placental function (Charalambous *et al.* 2003; Charalambous *et al.* 2010) and metabolism (Smith *et al.* 2007). In contrast, paternal *Grb10* is essentially restricted to the brain alone and impacts on behavioral phenotypes, including social dominance (Garfield *et al.* 2011). *Grb10* expression in the brain is neuronal, showing a pattern of expression in discrete brain regions that is shared with the maternally expressed imprinted gene *Nesp* (Plagge *et al.* 2005). This overlap in brain expression of maternal *Nesp* (encoding Nesp55) and paternal *Grb10* has led to speculation that these two genes may influence common adult behaviors, possibly in an opposite manner (Garfield *et al.* 2011; Dent and Isles 2014), which would be consistent with the prediction from the intra-genomic conflict theory of genomic imprinting evolution (Haig 2000).

Here, we address this question directly. We have shown that mice lacking maternal Nesp55 (*Nesp*^m/+^) make more impulsive choices (Dent *et al.* 2016). In contrast, here we show that *Grb10*^+/p^ mice make less impulsive choices relative to their littermate controls. Furthermore, there were no dissociable effects on a separate measure of impulsive action, completely matching the specificity of effects seen previously in *Nesp*^m/+^ mice (Dent *et al.* 2016). Our data provide new findings on *Grb10* function in brain and reveal opposite effects of manipulating *Grb10* and *Nesp* expression on discrete aspects of impulsive responding. Taken together these data are consistent with the conflict theory of the evolution of imprinted genes and as such provide the first direct evidence for intra-genomic conflict impacting adult behavior.

## RESULTS AND DISCUSSION

### *Grb10*^+/p^ mice make less impulsive choices

We used the delayed-reinforcement task to examine impulsive choice behavior (Isles *et al.* 2003), where subjects made a choice between receiving a small food reward after a short (1s) delay, or a larger reward after a longer delay (1, 8 or 16s). The extent to which subjects make impulsive choices is indexed by a preference for choosing the immediate but smaller reward, versus a larger delayed reward (delayed gratification). Increasing the delay to the larger reward within session decreased the likelihood of choosing that reward across all subjects (Figure 1A; main effect of DELAY, F_2,42_=9.20, *p*=0.001, partial η^2^=0.37). However, *Grb10*^+/p^ mice were significantly more tolerant of an increased delay to the larger reward than wild-type (WT) littermate mice (Figure 1A; main effect of GENOTYPE, F_1,21_=6.45, *p*=0.019, partial η^2^=0.25). However, whilst the data indicate that the experimental group diverges from wild-type increasingly as the delay increases, this effect failed to result in a significant interaction between GENOTYPE and DELAY (Figure 1A; F_2,42_=0.56, *p*=0.946, partial η^2^=0.003), due largely to a pre-existing group difference in the first 1s delay block.

**Figure 1.**
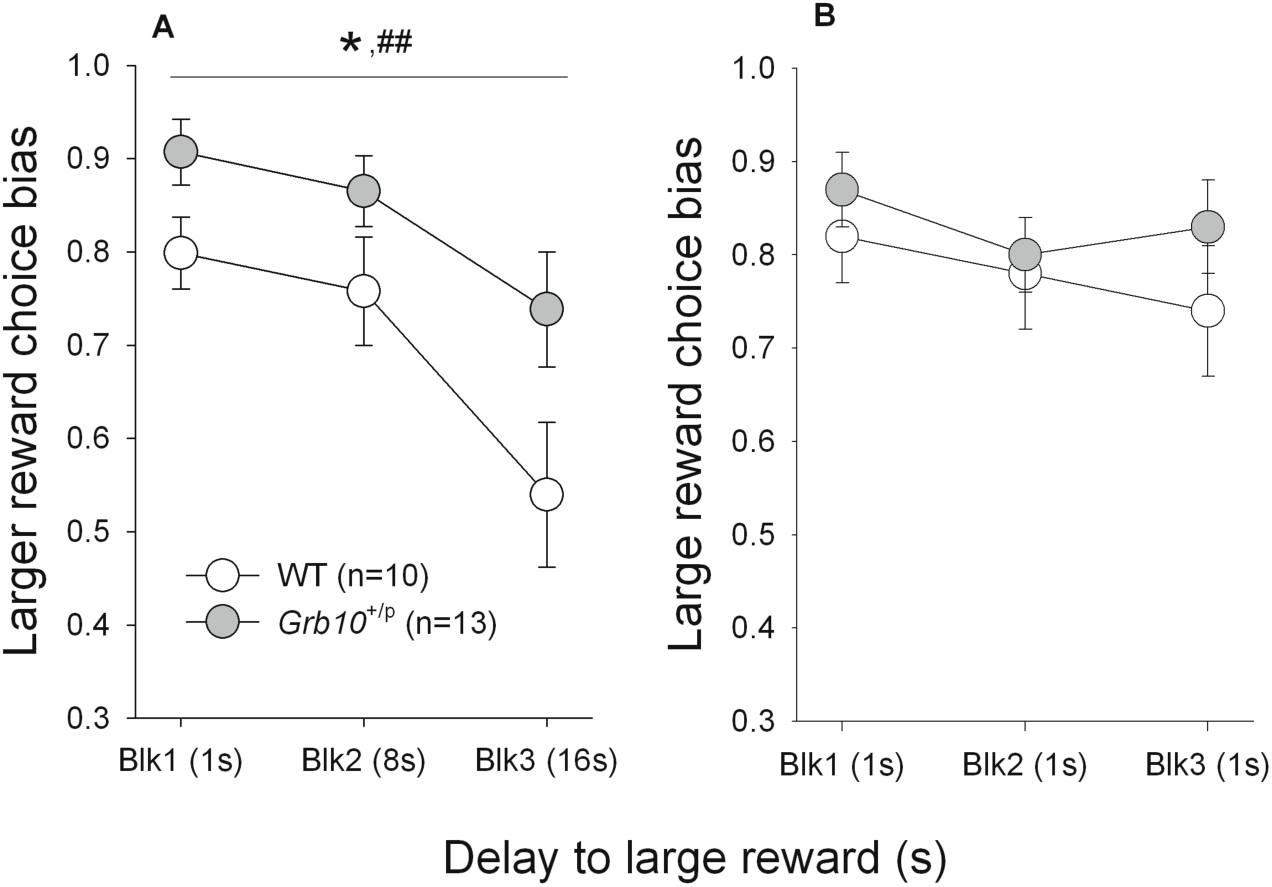
*Grb10*^+/p^ mice make less impulsive choices on a delayed-reinforcement task. Behavior of both *Grb10*^+/p^ and WT mice changed across session blocks (blk) with increasing delay, such that choice bias moved away from the response leading to the large reward towards the small reward, with increasing delay (**A**). However, there were systematic differences between the groups in their behavior, such that *Grb10*^+/p^ animals switched their choice to the small, less delayed reward less quickly than WT mice.

A standard control measure to assess the extent to which behavior is sensitive to delay is a task manipulation where any differential delay associated with the larger and smaller reward was equalized (1s) across all three blocks. Here, all subjects now demonstrated the expected preference for the larger reward throughout the session (Figure 1B; no main effect of DELAY, F_1.7,35.1_=1.88, *p*=0.17, partial η^2^=0.08); that is under these task conditions there were no differences in choice-bias between *Grb10*^+/p^ and WT mice (no main effect of GENOTYPE, F_1,21_=0.63, *p*=0.44, partial η^2^=0.03). Furthermore, there was no difference in the sessions taken to achieve a stable baseline in these conditions (WT 9 ±0.8; *Grb10*^+/p^ 8 ±0.5; *t*_21_=1.51, p=0.15).

The contrasting patterns of choice behavior between the *Grb10*^+/p^ and WT mice was also not due to any differences between the groups in terms of basic motivation to carry out the task as *Grb10*^+/p^ animals acquired the task at the same rate as WT littermates (sessions to last day of baseline; WT 50.4 ±1.1 SEM, *Grb10*^+/p^ 47.7 ±1.2 SEM *t*_21_=1.73, *p*=0.10). Additionally, variability between *Grb10*^+/p^ and WT mice was not related to differences in experiencing the information trial contingencies. In the ‘forced’ trials (where no choice was available), both *Grb10*^+/p^ and their WT littermates made equal responses to the large and small reward-related stimuli at all delays (no interaction GENOTYPE x DELAY x CHOICE, F_2,42_=0.44, *p*=0.649; data not shown). Finally, there was no difference between *Grb10*^+/p^ and WT mice on general measures of task performance (see Supplemental Information).

When the delay associated with the large and small rewards was equal (1s) throughout the session (**B**), choice bias was consistently high (large reward chosen approximately 80% of the time). Under these task conditions there were no differences in choice bias between *Grb10*^+/p^ and WT mice. Data shows mean ±SEM of three consecutive stable sessions; * represents P<0.05 main effect of GENOTYPE; ## represents P<0.01 main effect of DELAY.

In large part choice bias behavior in delayed reinforcement tasks is governed by three potentially interacting psychological processes; the perceived value of the rewards, and the perceived length and aversive nature of the delay (Ho *et al.* 1999). The difference in choice between *Grb10*^+/p^ and WT mice in the first block of the task (where delay for the larger and smaller rewards are both 1s) may suggest a contribution to behavior in the *Grb10*^+/p^ from a relatively more less pronounced generalized conditioned place aversion to the delayed larger reward response (see Isles *et al.* 2004). However, importantly, such an effect did not lead to an inflexible bias in responding as demonstrated by the continued sensitivity to delay, both within a baseline session and in response to the equal-delay probe manipulation. Therefore, whilst the between-group behavioral differences were clearly influenced to a large degree by a differential tolerance to delay the present data do not rule out influences due to aversion and reward perception

### *Grb10*^+/p^ mice show no difference in impulsive action

In contrast to impulsive choice, there was no difference between *Grb10*^+/p^ and WT mice on a measure of impulsive action. The stop-signal reaction time (SSRT) task measures the ability to stop an action once initiated by presenting a stop-signal (in this case an auditory tone) during a rapid response between two stimuli locations (the ‘go-response’). Correctly inhibiting the go response will earn reward in a ‘stop’ trial, whereas reward was also presented on completing a go-response in trials where no stop-signal was presented. Throughout the training stages of the SSRT task, all subjects showed show equivalent behavior in learning the task and *Grb10*^+/p^ and WT mice acquired the task at the same rate (sessions taken to complete the task, *Grb10*^+/p^: 42.4± 9.8, WT: 40.8±4.5, *t*_20_=0.54, *p*=0.59).

As expected (Humby *et al.* 2013; Davies *et al.* 2014; Dent *et al.* 2016), presenting the stop-signal progressively closer to the execution of the response (10, 40, 50, 60 and 90% into the individualised go response) led to systematic reductions in the ability to stop for all mice (Figure 2A; ANOVA, main effect of STOP, F_4,80_=24.37, *p*<0.001, partial η^2^=0.55). However, there were no differences in stopping efficiency between *Grb10*^+/p^ and WT mice (Figure 2A, main effect of GENOTYPE, F_1,20_=0.94, *p*=0.34, partial η^2^=0.05). The lack of genotype differences in stopping ability was further demonstrated by another measure of response inhibition, the speed of stopping or stop-signal reaction time (SSRT) derived in sessions where the subjects exhibited 50% correct stopping (Bari *et al.* 2009; Humby *et al.* 2013; Davies *et al.* 2014; Dent *et al.* 2016). Equivalent SSRTs were observed in both *Grb10*^+/p^ and their WT littermates (Figure 2B; *t*=17, *p*=0.87).

**Figure 2.**
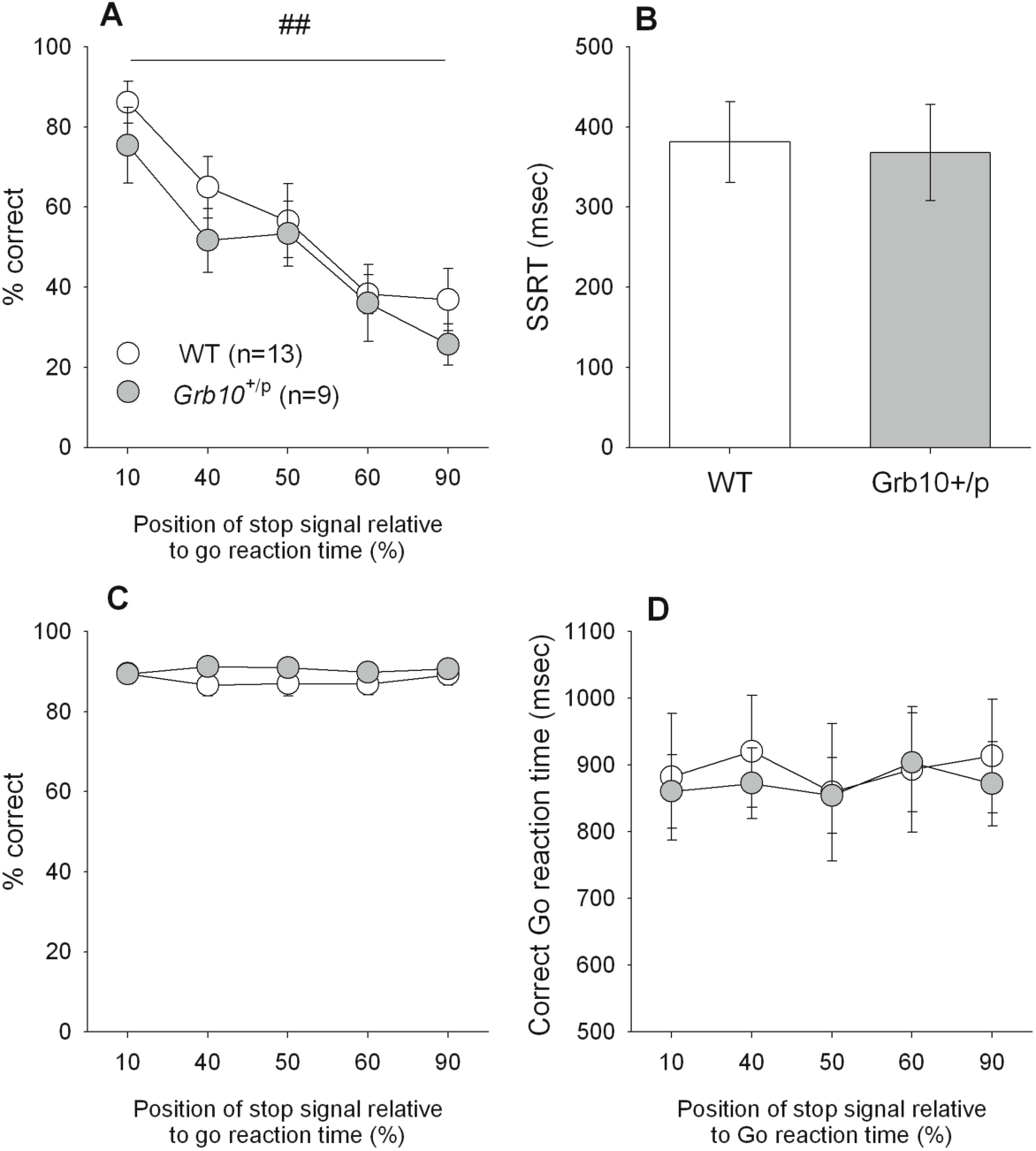
No difference beween *Grb10*^+/p^ and WT mice in performance on a stop-signal reaction time (SSRT) task. Both WT and *Grb10*^+/p^ mice showed an equivalent ability to perform the SSRT task showing the expected change in percentage correct responding during a ‘stop’ trial (**A**) as the position of the stop-signal was altered, but there were no differences between *Grb10*^+/p^ and WT mice. *Grb10*^+/p^ and WT mice also showed equivalent SSRTs at 50% correct stopping (**B**). There were no genotype differences for the Go-response for both groups of mice, in terms of percentage correct responding **(C)** or response speed (**D**). Data shows mean ±SEM; ## represents P<0.01 main effect of stop-signal position

There were also no differences in the Go-response between *Grb10*^+/p^ and WT mice, in terms of the amount (Figure 2C, main effect of GENOTYPE, F_1,20_=1.14, *P*=0.30) or speed (Fig. 2D, main effect of GENOTYPE, F_1,20_=0.03, *P*=0.86), of correct responding in Go trials. These parameters were not affected in sessions when the stop-signal position was moved from baseline (main effect of STOP-SIGNAL POSITION, F_4,80_=0.28, *P*=0.89 and F_4,80_=0.85, *P*=0.50, for the amount and speed of correct responding in Go trials, respectively). Additionally, there was no difference between *Grb10*^+/p^ and WT mice on general measures of task performance during individualized SSRT session (see Supplemental Information), indicating a high degree of stimulus control for both groups in the task.

### Opposite effects to maternal *Nesp*

The current data obtained from knocking out the paternal copy of *Grb10* mirror are essentially opposite to our previously published effects of knocking out the maternal copy of Nesp55 (Dent *et al.* 2016). That is, under identical task conditions, mice lacking maternal Nesp55 make more impulsive choices, whereas mice lacking paternal *Grb10* make fewer impulsive choices, relative to their littermate controls. These effects were highly specific in that they occurred in the absence of deficits in either model on responding in the SSRT task. The idea that these oppositely imprinted genes converge on discrete aspects of impulse control is further supported by their general overlapping expression patterns in the brain (Plagge *et al.* 2005; Garfield *et al.* 2011). In the present work we extend these neuroanatomical data in showing that Nesp55 and Grb10 are co-localised in cells in the locus coeruleus (Figure 3a-c), hypothalamus (Figure 3d-f), and the dorsal raphe nucleus (Figure 3g-i). Together, these studies indicate that impulsive choice behavior is a substrate for the action of genomic imprinting, and an extension of this idea is the speculation that an imbalance in expression of imprinted genes, such as *GRB10* or *NESP*, may also contribute to pathological conditions, such as pathological gambling, where choices become highly maladaptive. In this regard it should be noted that neither *GRB10* nor *NESP* have associated SNPs identified as significant variants in recent GWAS studies of gambling behaviour (Lang *et al.* 2016), or indeed delay discounting (Sanchez-Roige *et al.* 2018). However, it is worth noting also that genes with known roles in impulsive behaviour (eg *DAT, DRD1* or *DRD2*) were not associated either, suggesting that a lack of signal in GWAS does not necessarily diminish the suggestion of a role for *GRB10* and/or *NESP* in these disorders of impulse control.

**Figure 3.**
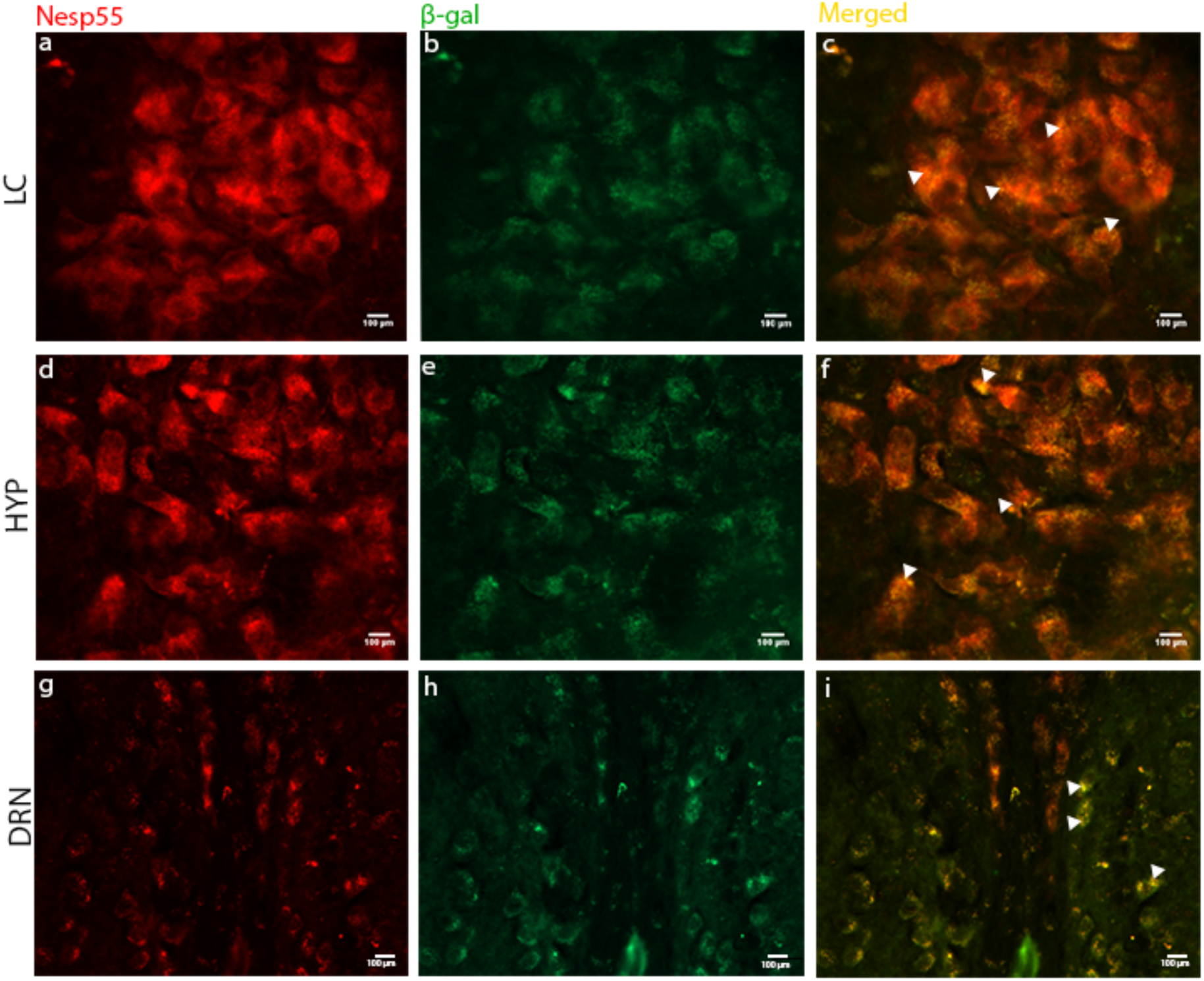
Dual-labelling immunofluorescence histochemistry of Nesp55 and Grb10 in coronal sections of adult brain. Sections were dual-labelled with antibodies against Nesp55 and β-gal, where the reporter gene LacZ is expressed in place of Grb10 in tissue from Grb10^+/p^ mice, and can be used to identify Grb10-positive cells. Images were viewed at different light intensities (568nm and 488nm for Nesp55 and b-gal, respectively), and were then merged to gauge cellular co-localisation of the two target proteins, depicted by white arrows in the merged figures. The majority of cells showed evidence for co-localisation: within the locus coeruleus (LC) (a-c), the hypothalamus (HYP) (d-f) and the dorsal raphe nuclei (DRN) (g-i). LC and HYP images at x40 magnification, DRN at x20.

We also suggest these data are consistent with the intra-genomic conflict theory of the evolution of imprinted genes, providing the first direct evidence for oppositional effects of imprinted genes on an adult behavior. In this context, the examples of Nesp55-and *Grb10*-null mice are particularly important, as these models are free from confounding effects on *in utero* or pre-weaning growth (Plagge *et al.* 2005; Garfield *et al.* 2011). Hence, this strengthens the argument that imprinted gene effects on adult behavior have adaptive significance and are not simply epiphenomena, that is, enduring into adulthood but resulting from the more familiar imprinted gene substrate of resource allocation between mother and offspring (Constancia *et al.* 2004).

Imprinted genes are thought to have evolved as a consequence of conflicting phenotypic “interests” between maternal and paternal genes that causes an escalating arms race in relation to allelic expression, eventually leading to silencing of one or other parental allele (Moore & Haig, 1991, Wilkins & Haig, 2003). Evidence of this parental conflict is seen in the contrasting action of maternal and paternal imprinted genes during *in utero* growth and early post-natal life (Haig, 2004). Although the present work is the first demonstration of opposing parental interests on behavior, the specificity of imprinting effects on impulsive choice behavior described here would fit with previous suggestions that maternal and paternal genomes may have a contrasting impact on decision-making in social animals (Haig 2000; Brandvain *et al.* 2011). Patterns of impulsive choice behavior with increasing delay, as measured by delay-discounting in the delayed-reinforcement task here, are often thought of as ‘irrational’, in that the shape of behavior is not consistent with simple models of optimization (such as expected utility maximization) (Schonberg *et al.* 2011). It may be that the so-called “parliament of the mind” (Haig 2000), caused by opposing parental genomes pulling impulsive choice in different directions, is an additional factor that should be taken into consideration in the context of more complex models of apparently ‘irrational’ behaviors (Fawcett *et al.* 2014).

## MATERIALS AND METHODS

### Animals

All procedures were conducted in accordance with the UK Animals (Scientific Procedures) Act 1986. Subjects were male mice, and were 4 months old at the start of testing, which was completed after 6-months. Standard laboratory chow was available *ad libitum*, but during the experiment water was restricted to 2h access per day. This regime maintained the subjects at ≈90% of free-feeding body weight. Due to potentially confounding phenotypes associated with *Grb10*^m/+^ mice, comparisons of *Grb10*^+/p^ were made with wild-type (WT) littermate controls derived from 8 separate litters. The *Grb10* null line was maintained on an F1-hybrid (C57/Bl6 X CBA) background. The same cohort was used throughout testing but a number of animals were lost (due to death or inability to perform training stages) as testing progressed.

### Behavioral testing

#### Operant testing apparatus

All the sessions of the delayed reinforcement task and SSRT task were performed in 9-hole operant chambers (Cambridge Cognition Ltd, U.K) modified for use in mice, as described previously (Humby *et al.* 1999; Isles *et al.* 2003; Humby *et al.* 2013). For the delayed reinforcement task holes 3, 5 and 7 were open; whereas only holes 4 and 6 were open for the SSRT task. The mice were presented with a visual stimuli (light) recessed into the holes and were trained to respond to this stimulus with a nose-poke as recorded by infra-red beams spanning the hole. Reward was presented in a recessed compartment on the wall opposite to the nose-poke/stimulus array. The control of the stimuli and recording of the responses were managed by an Acorn Archimedes computer with additional interfacing by ARACHNID (Cambridge Cognition Ltd). For all operant testing, animals were maintained on a restricted water access schedule, water provided for two hours immediately after testing.

#### Delayed reinforcement task

Details of the shaping procedures and basic aspects of the delayed-reinforcement task itself can be found elsewhere (Isles *et al.* 2003). Briefly, the task comprised of three sequential blocks of 12 trials, with each trial consisting of an initial nose-poke to the centrally located stimulus, followed by a second nose-poke to either the left or right apertures. Trials 1-4 in any block were ‘forced’ information trials, where the initial nose-poke resulted in presentation of only one of the two choice options. This measure was designed to provide the subjects with prior notice of the extent of any delay associated with choosing the large reward. In the remaining 8 trials of each block, designated as ‘choice’ trials, the initial centre nose-poke led to the option of a second nose-poke response to either the left or right apertures. One response resulted in the delivery of a large reward (50μL 10% solution of condensed milk; Nestle Ltd, UK), and the other in the delivery of small reward (25μL 10% solution of condensed milk). The response contingencies were kept constant for each mouse, but were counterbalanced between subjects. In block 1 both responses led to the delivery of reward after a 1s delay. In blocks 2 and 3 increasing delays were introduced between the response and the delivery of the large reward (8s, 16s, respectively) whereas the delay between response and delivery of the small reward was fixed at 1s. As a probe to test the effect of the delays on behavior, sessions were conducted where the delay associated with the large reward was fixed at 1s, equivalent to that associated with the small reward, throughout all three blocks of the session.

The bias in choice of the larger reward at each block (whereby always choosing the large reward=1; never choosing the large reward=0) was the main measure used to determine impulsive responding. Additional measurements that related to general motoric competence and motivation within the task were also monitored, including the ‘Start’ and ‘Choice’ latencies, the time taken to initiate a trial and the time taken to make a choice once a trial was initiated, respectively. Also measured were the number of ‘Non-started’ (no initial, central nose-poke) and ‘Omitted’ (no secondary, choice nose-poke, following central nose-poke initiating trial) trials.

#### Stop-signal reaction time (SSRT) task

Details of the shaping procedures and basic aspects of the main SSRT task can also be found elsewhere (Humby *et al.* 2013). The SSRT task itself consisted of sessions of 100 trials, which involved both ‘go’ and ‘stop’ trials. Go trials consisted of rapid double nose-pokes (a ‘go’ response) between two separate stimuli locations, which were rewarded with reinforcement (22ul, 10% solution of condensed milk, Nestle Ltd, UK). 20% of trials were stop trials, pseudo-randomly distributed throughout each session, where a stop-signal (65db white noise for 0.3s) was presented between the first and second nose-poke responses. The aim of the stop-signal was to inhibit (‘Stop’) the mouse from making the second (‘Go’) nose-poke, and then wait for the reward. Failure to refrain from making this pre-potent response was punished by the absence of reward and 5 second time out (chamber light on). At baseline, the stop-signals were presented concurrently with the initial nose-poke response. To maintain high levels of performance of both go and stop responding, the go stimulus duration and wait period to reward delivery in a stop-signal trial were determined individually for each subject. To assess the ability to stop once an action had been initiated, sessions were implemented in which the onset of the stop-signal was presented at different positions within the individualised go response of each mouse. Thus, the stop-signal was pseudo-randomly presented 10, 40, 50, 60 and 90% from the onset of the go response of each subject, with the assumption that stopping would be more difficult the closer the stop-signal presentation was to the termination of the go response.

The amount of correct stopping in stop-signal trials and the SSRT were the main measures of impulsive responding in this task. The SSRT was calculated by determining the 50% stopping ability for each subject from the range of sessions in which the stop-signal onset was varied from baseline (full details of this calculation can be found in the Supporting Information and (Davies *et al.* 2014)). The proportion of correct go-responses, and latency to respond, were also assessed. Additional measurements that related to general motoric competence and motivation within the task were also monitored, including the “initiation” and “magazine” latencies, the time taken to initiate a trial and the time taken to collect the reward. Also measured was the number of trials completed for any given session.

### Immunohistochemistry

Dual-labelling immunofluorescence analysis of Nesp55 co-localisation with Grb10 was carried out on brain sections from *Grb10*^*+/p*^ mice in order to stain for the Lac-Z reporter gene, which is expressed in place of *Grb10* and has been used as a faithful proxy for Grb10 expression previously (Garfield *et al.* 2011). *Grb10*^+/p^ mice were transcardially perfused using 10% formalin (Sigma-Aldrich, UK) and whole brains dissected, post-fixed and equilibrated with 30% sucrose in PBS. Brains were sectioned into 40μm coronal slices using a freezing microtome. Sections were washed three times for 10 min each in 0.1% PBS before being incubated for 15 min in 0.3 M glycine in 0.1% PBS at room temperature, to neutralise endogenous aldehyde groups. Sections were washed, as before, in 0.1% PBS and then incubated at room temperature for 1 hour in 10% blocking solution; 0.5% BSA (BB International, Cardiff, UK), 0.5% Triton X-100 (v/v, Sigma Aldrich) in 0.1% PBS. Sections were then transferred to a 1% blocking solution containing a β-galactosidase (β-gal) specific antibody (Abcam, U.K.), used at a 1:1000 dilution; and a Nesp55 primary antibody (1:1000). The Nesp-55 primary antibody was generated in house, and is a rabbit anti-Nesp55 polycolonal antibody recognising the free terminal end (GAIPIRRH) of Nesp55. It has been successfully characterised and used previously (Ischia *et al.* 1997). Sections were incubated overnight at 4°C whilst gently shaking. Sections were then washed three times for 10 minutes in 0.1% PBS and then incubated with the appropriate fluorescent secondary antibodies (Alexa Fluor; Life technologies) (1:1000) in 1% blocking solution in the dark at room temperature for 2 hours, whilst gently shaking. Sections were then washed in 0.1% PBS as before (in the dark) and transferred to polysine coated slides and allowed to dry over-night in a dark dust-free environment. The mounted slides were then dehydrated through a process of incubation in a rising concentration of alcohol, followed by xylene, then cover-slipped and sealed using DPX (Raymond Lamb DPX), and allowed to dry over-night. To control for non-specific binding of the secondary antibodies, secondary-only negative controls were carried out alongside all experiments. Immunofluorescence slides were viewed and images captured using an upright fluorescence microscope (Leica DM5000 B). Dual-labelled immunofluorescence images were acquired through separate channels for different wavelengths (488 and 568 nm) then subsequently merged using ImageJ (Image>colour>merge channels).

### Data analysis and statistics

All behavioral data were analysed using SPSS 20 (SPSS, USA). Data were assessed for normality and then analysed by Student’s t-test or mixed ANOVA, with between-subjects factors of GENOTYPE (*Grb10*^+/p^ vs. WT), and within-subject factors DELAY (1s,8s,16s, or 1s,1s,1s), CHOICE (choice of large or small reward during forced trials of the delayed reinforcement task), and STOP-SIGNAL POSITION (position of stop-signal relative to individualized Go-response). For repeated-measures analyses, Mauchly’s test of sphericity of the covariance matrix was applied; significant violations from the assumption of sphericity were subject to the Huynh–Feldt correction to allow more conservative comparisons through adjusted degrees of freedom. Variables that were expressed as a percentage (Delayed reinforcement, choice bias; SSRT, % correct) were subjected to an arcsine transformation in order to limit the effect of an artificially imposed ceiling. All significance tests were performed at alpha level of 0.05. Effect sizes (partial η^2^) were reported for the main measures in all tasks.

### Data statement

Supplementary 1 contains additional control measures in the Delay discounting task (Figure 1 and Table1) and SSRT task (Figure 2); further Methodological detail relating to SSRT measure. Individual animal data for the delayed reinforcement task can be found in Appendix S1 (Excel spread sheet). Individual animal data for the SSRT task can be found in Appendix S2 (Excel spread sheet). All other datasets in the current study are available from the corresponding author on reasonable request.

## ACKNOWLEDGMENTS

This work was funded by a Leverhulme Trust project grant (F/00 407/BF) to ARI and JFW, which supported CLD. The work was also funded by the MRC Centre for Neuropsychiatric Genetics and Genomics (G0801418), of which ARI, TH and LSW are members, and which also supported KL via a PhD studentship.

