## Supplementary Materials for "Impulsive choice in mice lacking paternal expression of *Grb10* suggests intra-genomic conflict in behavior"

**SUPPLEMENTARY MATERIALS AND METHODS**

**Calculating the Stop Signal Reaction Time**

- Correct go reaction times were determined directly, however the stop-signal reaction time (SSRT) had to be derived from the distribution of correct go reaction times and the proportion of correctly stopped trials. SSRTs in the task were estimated employing the standard procedure previously described (Logan *et al.* 1984), using data from where the proportion of correct stop responses was ~50%. For each subject, data from the sessions in which the stop-signal positions were varied relative to the individualised go reaction time, were ranked by the proportion of correct stop responses, and data from sessions in which this value was between 40% and 60% (i.e. 50%±10%) were averaged. The latency of stopping as defined by the SSRT was derived from the distribution of correct go reaction times and the proportion of correctly stopped trials as previously described (Logan 1994; Eagle and Robbins 2003). Hence, for each of the sessions determined above, the correct go reaction times were rank ordered from smallest to largest and the n^th^ value found, where n is the rank order position based on the proportion of failing to stop correctly in stop trials was corrected for the occurrence of omitted go trials. Although omitted go trials were a rare occurrence, this correction was implemented based on the rationale that they could alter the observed inhibition function and affect determination of the n^th^ correct go reaction time value and hence the final SSRT (Tannock *et al.* 1989; Solanto *et al.* 2001; Eagle and Robbins 2003). To determine the SSRT, the time the stop-signal was presented (i.e. ‘mean correct go reaction time’ x ‘% mean stop-signal position’) was subtracted from the n^th^ correct go reaction time value.
- **SUPPLEMENTARY RESULTS**

**Additional measures in the delayed reinforcement task**

In addition to choice, a number of other measures of basic behaviour within the delayed reinforcement task were obtained. These relate to general motoric competence and motivation within the task. As expected, *Grb10*^+/p^ and wild-type (WT) littermates demonstrated a general main effect of DELAY on all of these measures as follows: Start (Figure S1a; F_1.4,26.8_=11.76, *p*=0.001) and choice latencies (Figure S1b; F_1.8,33.5_=31.5, *p*<0.001); and non-started trials (Figure S1c; F_1.6,30_=21.95, *p*<0.001) and omitted trials (F_1.3,24.6_=12.48, *p*<0.001). In all of these measures *Grb10*^+/p^ and WT littermate mice showed a high degree of stimulus control, with no GENOTYPE differences or interactions between GENOTYPE and DELAY (see Table S1). Data for Start latency, Choice latency and non-started trials can be found in Figure S1. Omitted trials occurred very infrequently (Grand mean 0.21, ±0.05 SE), and are not shown.

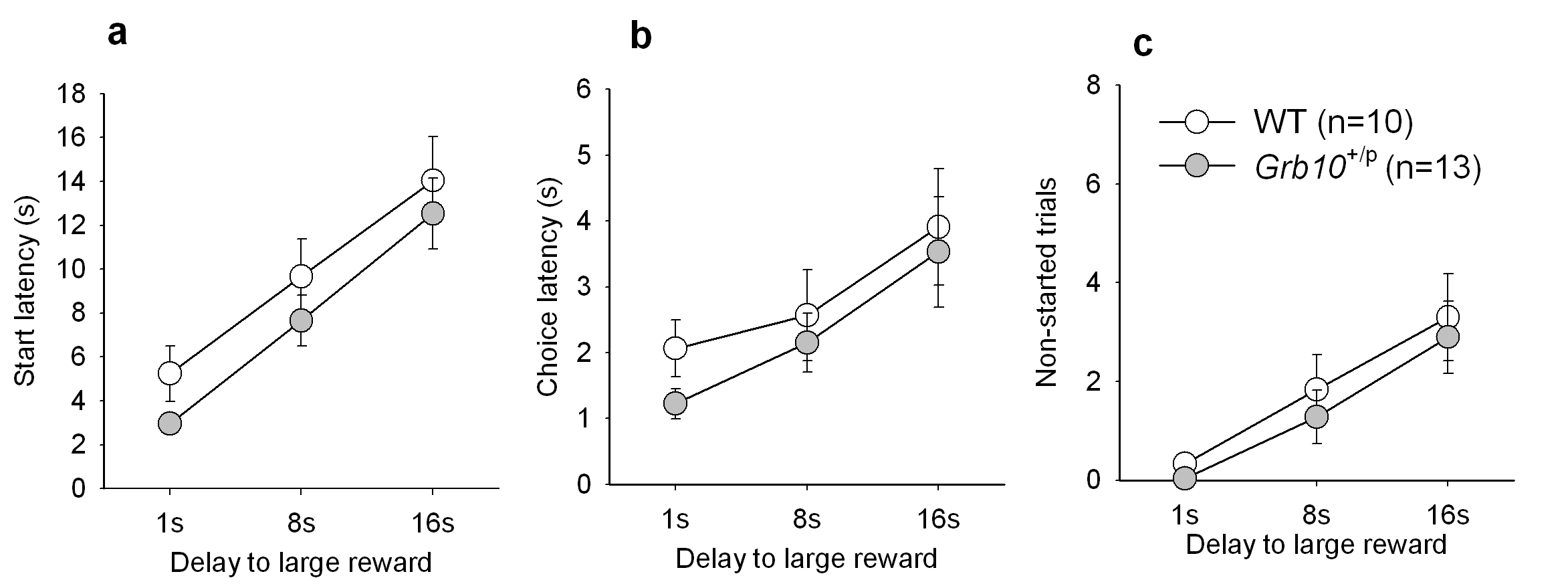

**Figure S1 Additional behavioural measures of performance of *Grb10*^+/p^ and WT mice in the delayed reinforcement task.** **a** Start latency; **b** Choice latency; **c** Non-started trials; **d** Panel pushes. All data represent means ±s.e.m.

| **Measure** | Main effect of GENOTYPE | | Interaction: GENOTYPE and DELAY | |
| --- | --- | --- | --- | --- |
|  | F | *p* | F | *p* |
| Start latency | 2.63 | 0.12 | 0.58 | 0.51 |
| Choice latency | 0.45 | 0.51 | 0.10 | 0.88 |
| Non-started trials | 0.35 | 0.56 | 0.06 | 0.94 |
| Omitted trials | 0.01 | 0.93 | 0.62 | 0.48 |
| **Table S1 Statistical values for additional measures of performance of *Grb10*^+/p^ and WT littermate controls within the delayed-reinforcement task.** Analysis of main effect of GENOTYPE, and interaction between GENOTYPE and DELAY. | | | | |

**Additional measures in the SSRT task**

In addition to the main measures, a number of other measures of basic behavior within the SSRT task were obtained (Figure S3). There were no differences between *Grb10*^+/p^ and WT littermates in the number of trials completed (*t*_20_=0.01, *p*=0.99), latency to initiate a trial (*t*_20_=-0.54, *p*=0.60) and magazine latency (*t*_20_=0.99, *p*=0.33).

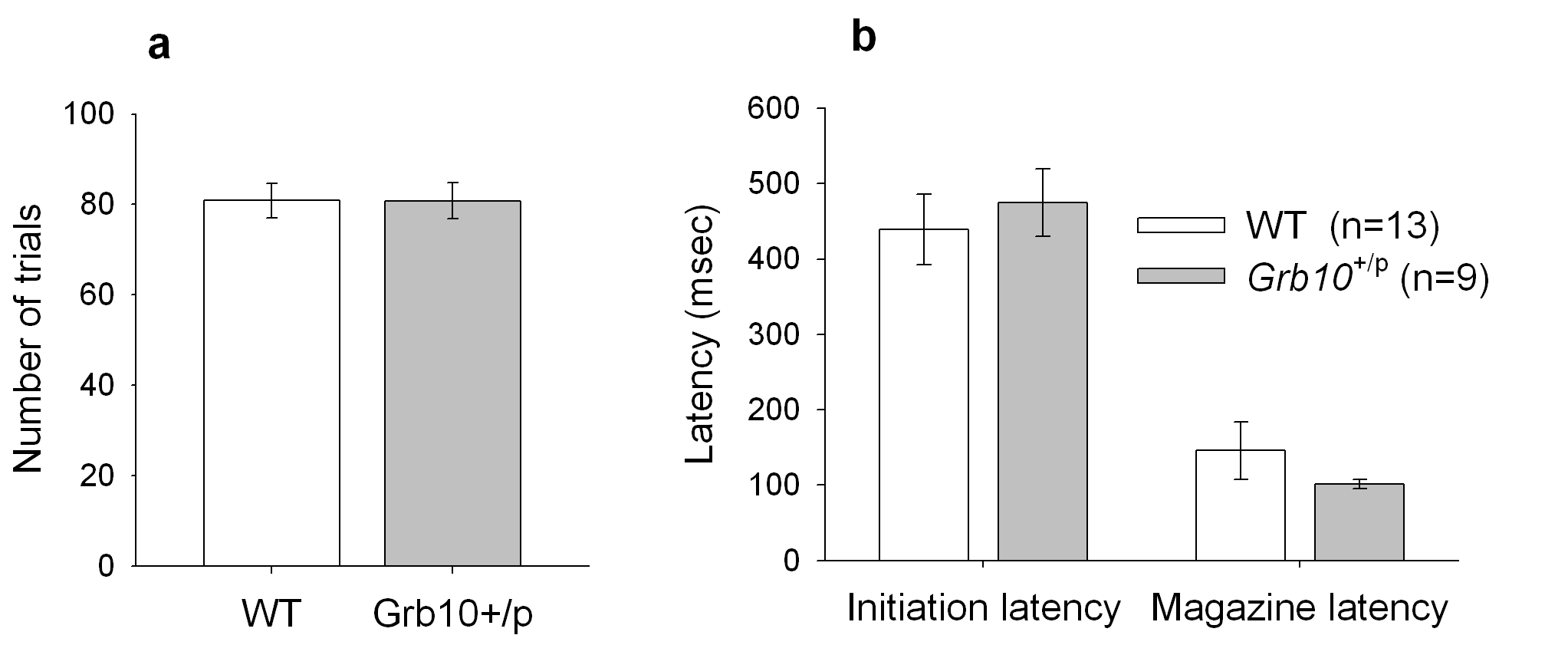

**Figure S2 Additional behavioural measures of performance of *Grb10*^+/p^ and WT mice in the SSRT task.** **a** Number of trials in a session with individualised SSRT (when the stop cue is presented 50% into the individual’s Go-reaction time); **b** Initiation and magazine latency in a session with individualised SSRT. All data represent means ±s.e.m.
